# Antibody-mediated depletion of protein variant expression in living cells (Protein interference)

**DOI:** 10.1101/119305

**Authors:** Vijay K. Ulaganathan, Axel Ullrich

## Abstract

A significant development in the field of human biology is the revelation of millions of unannotated protein sequence variants emerging from the several human genotyping and genome sequencing initiatives. This presents unique opportunities as well as confounding challenges in our understanding of how molecular signalling outcomes vary among individuals in the general population. As a result the conventional ‘one drug fits all’ lines of approach in the drug discovery process is becoming obsolete. However, an innovative genotype-specific approach targeting protein sequence variants instead of a reference protein target is currently lacking. In this short communication we report a remarkable observation of antibody-mediated knockdown of intracellular protein expression. This suggests allele-specific inhibition of protein-variant expression can be achieved by intracellular delivery of lipid conjugated linear epitope-specific monoclonal antibodies. The results presented here demonstrate novel opportunities for interrogating the protein coding variations in the human genomes and new therapeutic strategies for the inhibition of pathogenic protein variants in a genotype-centric manner.

## Introduction

With over five decades after the discovery of DNA double helix and over a decade after the complete sequencing of human genome we are still faced with the challenge of effectively dealing with aberrant and mutated proteins as underlying cause of many diseases. Furthermore new challenges arise with the ever-increasing knowledge of genomic variations prevalent among human populations. This calls on for a serious rethinking on the existing conventional strategy of drug discovery process based on “one drug fits all”. Recently, the prospect of identifying the genetic basis of diseases in the current genomic era has enormously increased. However, the big question of how to tackle specifically these disease causing protein variants without affecting the normal healthy alleles has remained unanswered. Small molecule chemical inhibitors purported to inhibit such mutated protein targets, often are binding and inhibiting multiple other proteins as well as wild type protein causing undesirable side effects. There are some positive anticipation from RNA interference and more recently the genome editing CRISPR technologies for knocking down specific proteins (Howard, 2003; Marraffini & Sontheimer, 2010). But these techniques have their limitation in not being able to distinguish protein variants differing in single amino acid change. While the recent genomic revolution has uncovered many patient specific targets in complex diseases such as cancer, the pace of discovering personalized solutions is still lagging behind (Drews, 2000; Roses, 2004).

Here we report a phenomenon akin to RNA interference, we call it ‘**Protein interference**’ which can be exploited for knocking down protein variants of interest to a specificity single amino acid or allelic difference. This new method offers not only a valuable tool for biological research but also paves way for a novel therapeutic approach to target proteins in allele specific manner.

## Results

Expression of protein variants in living cells were inhibited when high affinity monoclonal antibody raised against the linear peptide sequence of the antigen was delivered intracellularly. We show here that the levels of exogenously expressed green fluorescent protein (GFP) were significantly suppressed by the intracellular delivery of monoclonal antibodies raised against GFP (**Figure. 1**). Monoclonal antibodies (mAbs) raised against post-translational modifications such as phosphorylated tyrosine 4G10 did not deplete the levels of GFP. Interestingly, sequence specific downregulation of endogenous protein variants occurred when point-mutation specific mAbs were transfected to cells of different genotypes. Depletion of germline variants encoded by the single nucleotide polymorphism (SNP) rs351855, namely FGFR4 p.Gly388Arg (humans) and FGFR4 p.Gly385Arg (mouse) was observed when point-mutation specific mAbs were transfected to human and mouse cells of different genotypes respectively (**Figure. 2** and **Figure. 3**). The decreased levels of protein variant were verified by the abolishment of gain of function phenotype by the rs351855 SNP (Ulaganathan et al, 2015) in the germline genome. Interestingly, using this approach we succeeded in depleting challenging somatic variants namely BRAF p.V600E (**Figure. 4**) and KRAS p.G12V (**Figure. 5**). Furthermore, target specificity could be demonstrating by performing antibody transfection experiments under conditions of co-cultivation of genetically labelled MEFs stably expressing somatic variants namely BRAF p.V600E (GFP positive) and KRAS p.G12V (RFP positive) (**Figure. 6** and **Figure. 7**).

Preliminary experiments comprising of polycistronic reporter constructs (**Figure. 8**) suggests antibody mediated depletion of protein expression may involve RNA degradation. For instance, inhibition of GFP expression when HA epitope or KRAS p.G12V epitope was targeted (**Figure. 9**) suggests RNA degradation or translational inhibition. Based on these observations, we hypothesize the mechanism may be due to stalling of translation apparatus of protein biosynthesis caused by sequence specific interaction of antibody to polysomal polypeptide during biosynthesis. Any interference in polypeptide extension or entry of tRNA-amino acid during the protein biosynthesis process is subjected to translational quality control mechanisms by No-Go Decay (NGD) or Nonsense-mediated Decay (NMD) resulting in shutting down protein synthesis which is the most likely explanation for our results here (Lykke-Andersen & Bennett, 2014). This potentially represents a versatile approach to block protein variant of interest in a specific manner within a living cell for therapeutic or scientific research purposes.

**Figure 1.**
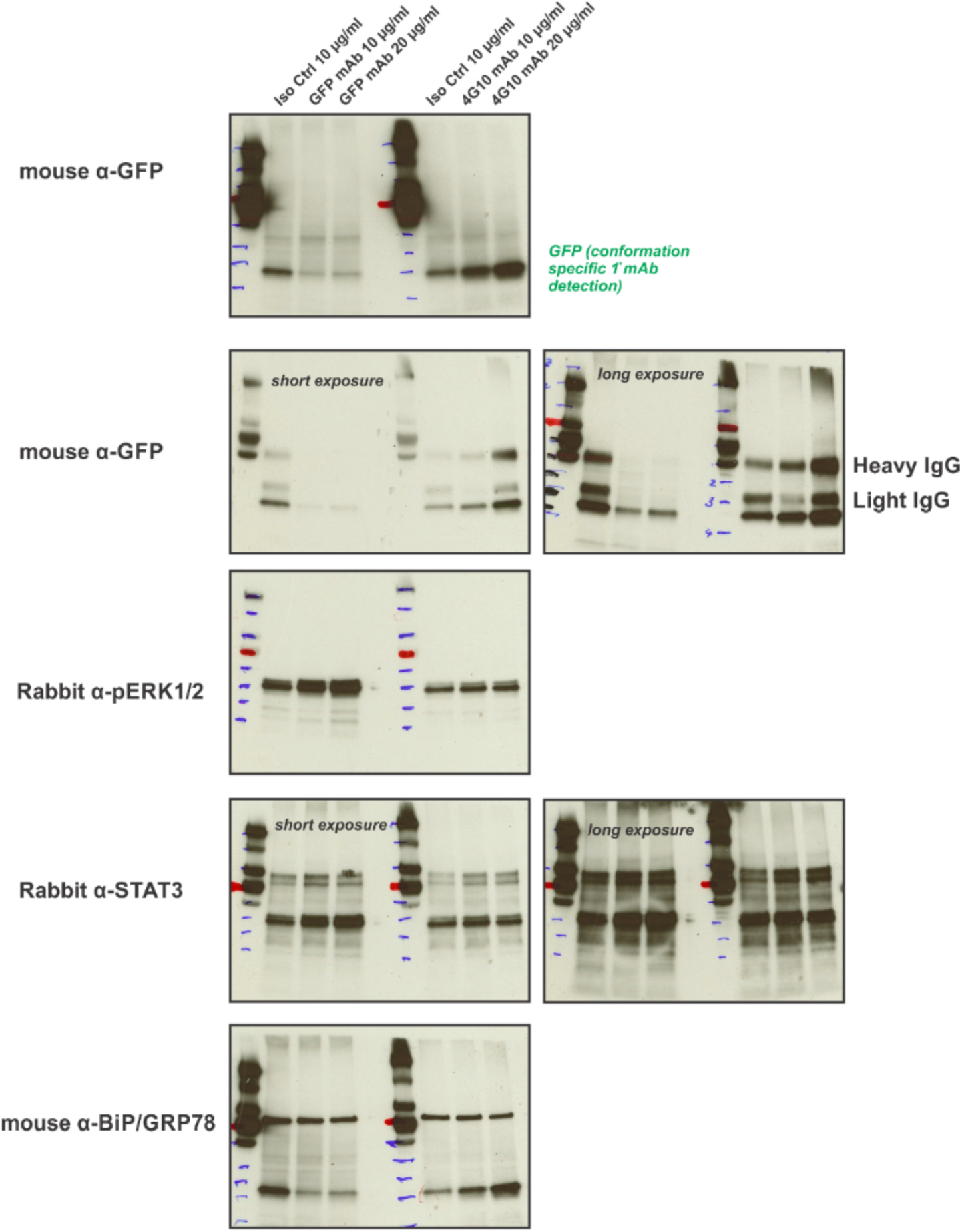
Specific depletion of exogenously expressed target protein. Indicated home-made monoclonal antibodies were delivered into cell lines expressing GFP under stably integrated CMV promoter using Pulsin (Polyplus) protein transfection reagent at a ratio of 1:2. Two days after protein transfection immonoblot analysis was performed using conformation specific secondary antibody that does not detect intracellular delivered IgG (first image) or conventional secondary antibodies. GFP expression was substantially depleted when immunoglobulins binding to GFP epitope was delivered whereas HA tag specific mAbs that did not bind to GFP target unaltered its expression. On the other hand phosphotyrosine specific mAb specific against post-translational modification instead of primary amino acid sequence did not result in downregulation rather resulted in an increased expression of GFP.

**Figure 2.**
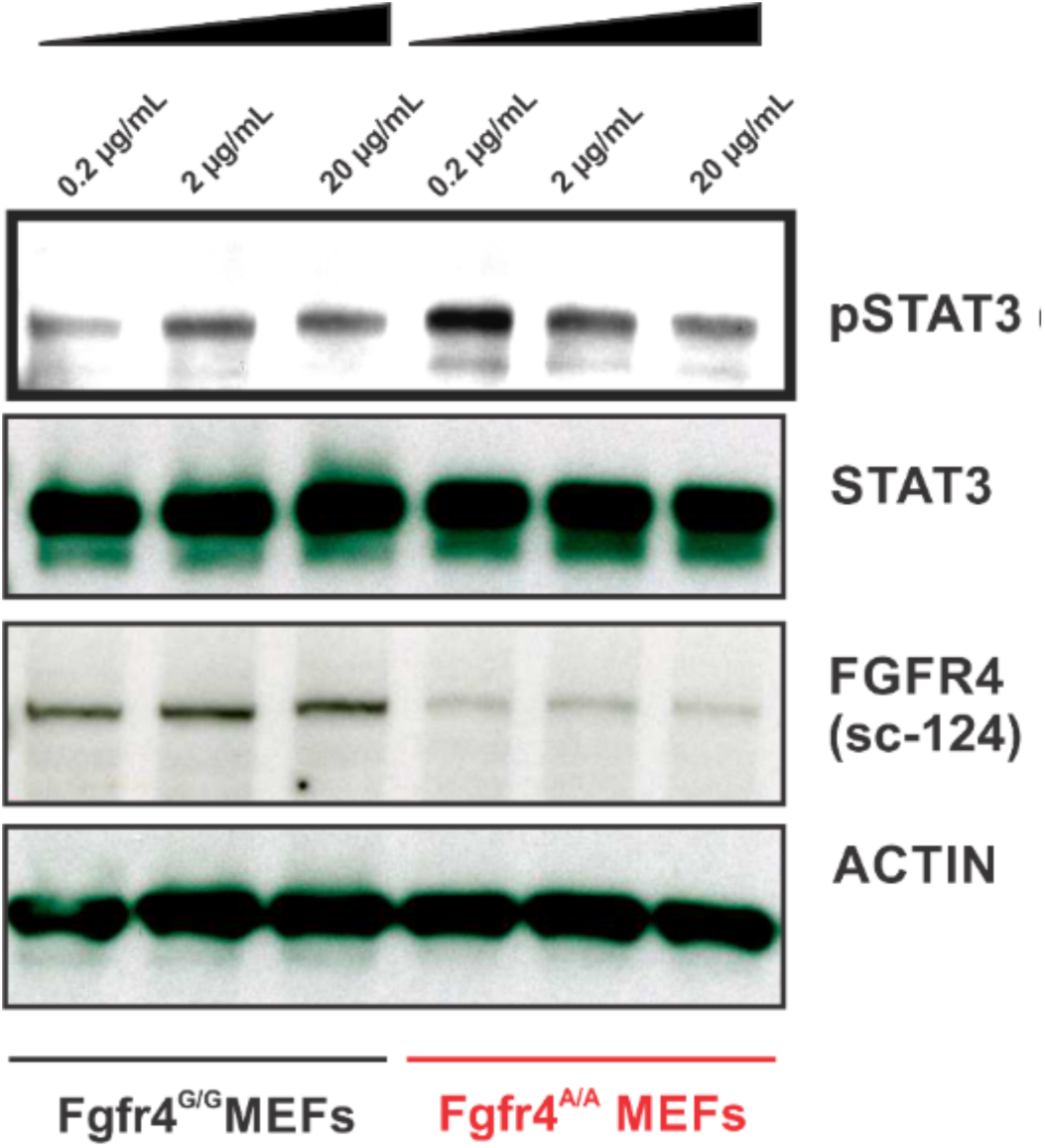
Precise depletion of *Fgfr4* germline variant (FGFR4 p. G385R) in mouse cells. FGFR4 Arg385 Point mutation specific mAb (custom made) were delivered into knock-in mouse embryonic fibroblasts derived from E13.5 mouse embryos of either genotypes (*Fgfr4^G/G^* genotype encode FGFR4 p.Gly385 protein & *Fgfr4^A/A^* genotype encode FGFR4 p.Arg385 protein; Entire genome of the two genotypes of mice are otherwise identical) at indicated amounts using Pulsin (Polyplus Inc) transfection reagent (1:2 ratio). Two days after mAb transfection expression levels of FGFR4 was assessed by immunoblot analyses. FGFR4 Arg385 is a gain of function mutation in the genome that causes increased basal expression of phosphorylated STAT3 (Y705)(Ulaganathan et al, 2015). Therefore, effect of pSTAT3 (Y705) was measured to ascertain the functional consequences of depletion of FGFR4 Arg385 variant as an indicator for therapeutic effect. Under identical transfection and growth conditions for 48 hrs only FGFR4 p.Arg385 protein variant was depleted thereby returning the levels of pSTAT3 (Y705) to normal level or that of wild type variant cells.

**Figure 3.**
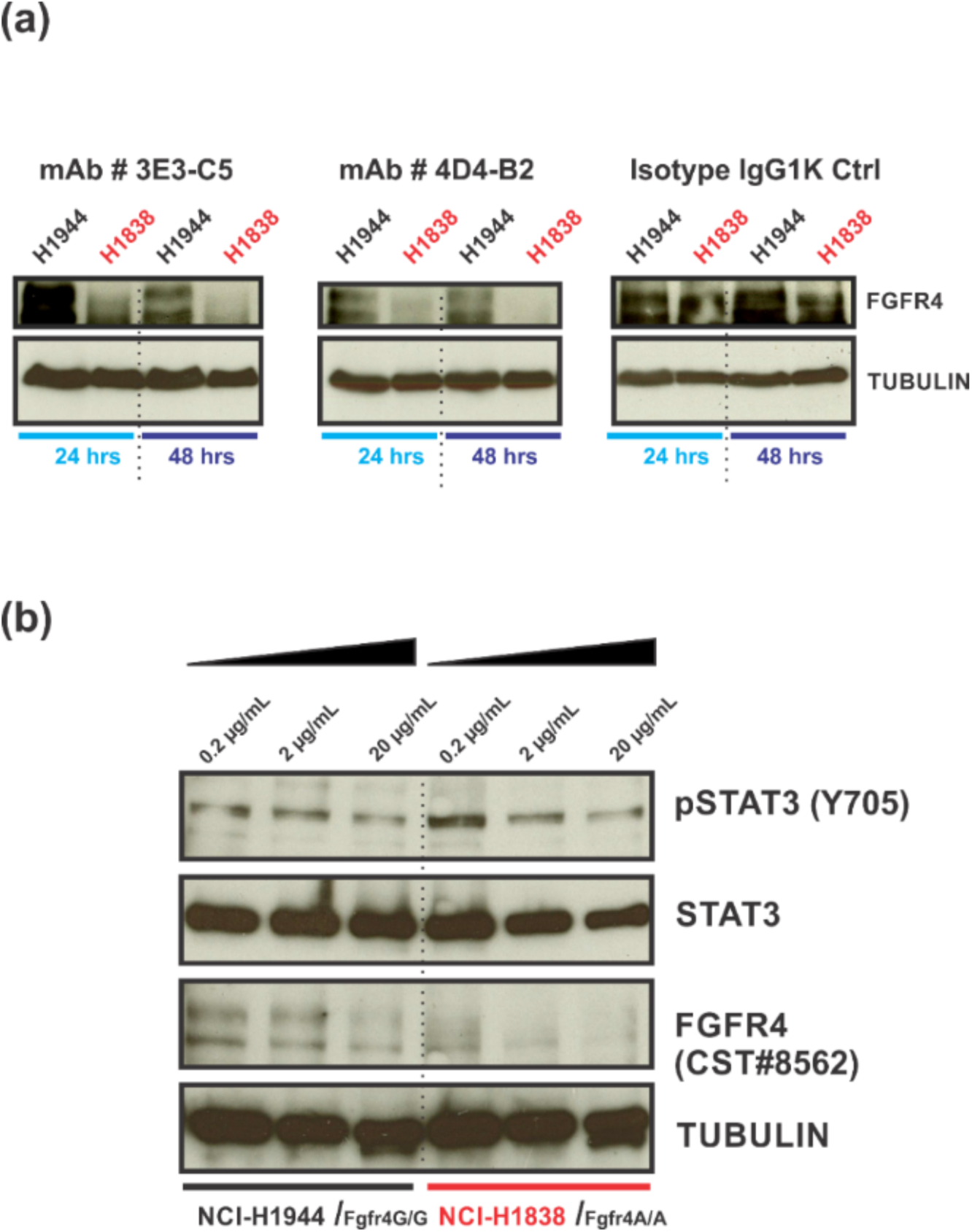
Precise depletion of *FGFR4* germline variant (FGFR4 p. G388R) in human cells. (a) FGFR4 Arg388 Point mutation specific mAb (custom made) clones were delivered into a human lung cancer cell lines (NCI H1838) expressing FGFR4 p. 388Arg protein variant encoded by the *FGFR4* germline gene variant *FGFR4* p.G388R (SNP ID: rs351855) and NCI H1944 that expresses wild type FGFR4 p.388Gly protein variant. (b) *FGFR4* p.G388R is a gain of function germline mutation in the genome that causes increased basal expression of phosphorylated STAT3 (Y705) in all tissues where *FGFR4* is expressed (Ulaganathan et al, 2015). Therefore, effect of pSTAT3 (Y705) was measured to ascertain the functional consequences of depletion of FGFR4 Arg388 variant to serve as an indicator for therapeutic effect. Under identical transfection and growth conditions for two days, only FGFR4 p.Arg388 protein variant was depleted subsequently resulting in returning the levels of pSTAT3 (Y705) to normal level.

**Figure 4.**
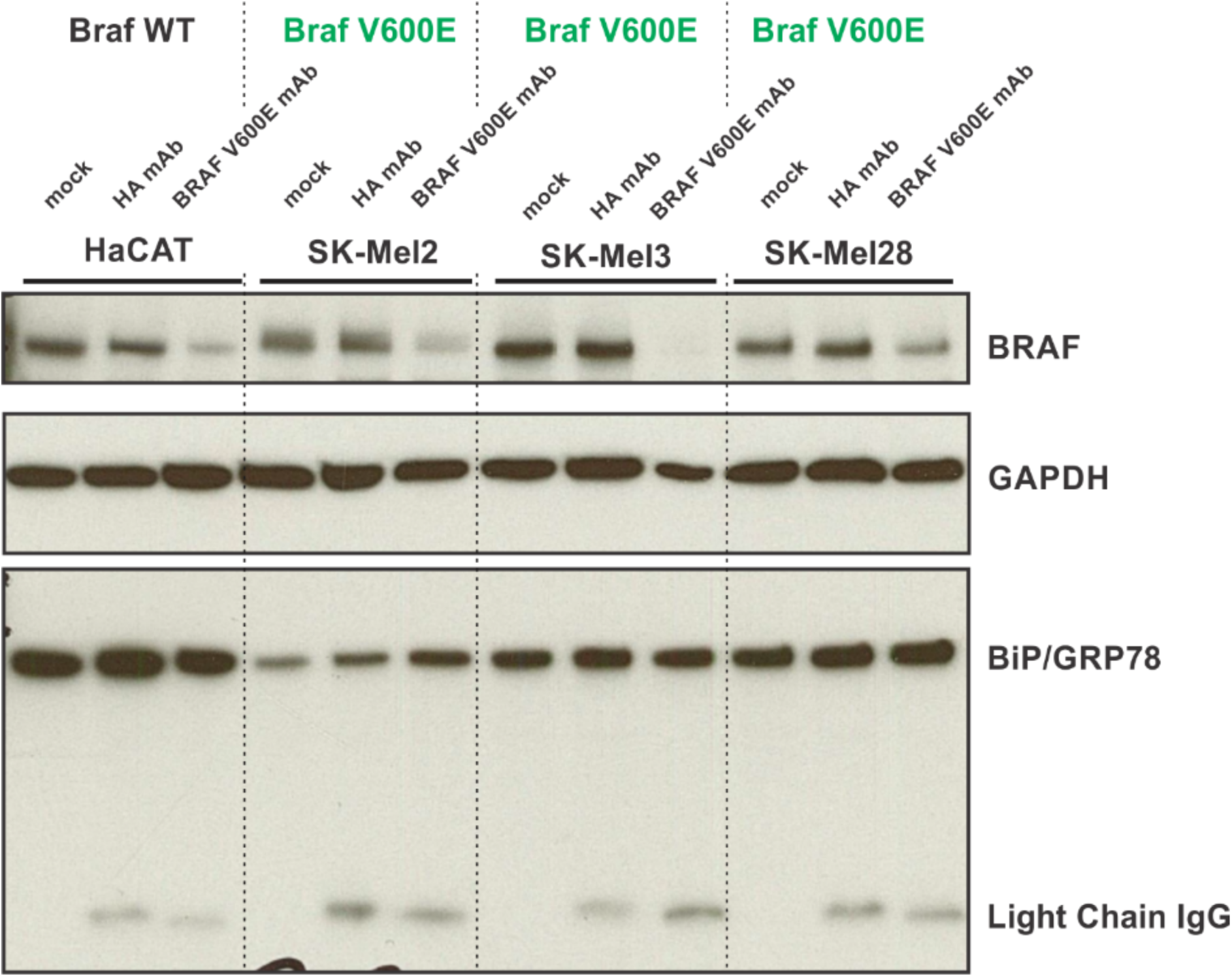
Specific depletion of *BRAF* somatic variant (BRAF p.V600E) in human skin cancer cells. BRAF 600E point mutation specific mAbs (New East Biosciences, preservative-free formulation) (10 μg) was delivered into human skin cancer cell lines (SK-Mel2, SK-Mel3, SK-Mel28) using ProJECT protein transfection reagent (Life Technologies) at 1:1 ratio. Non-malignant HaCAT cell line which express wild type BRAF gene was used as a control. Two days after transfection cell lysates were analysed by immunoblot for expression levels of BRAF. An apparent reduction in expression of BRAF protein was evident only with mutation specific mAbs but not with anti-HA mAbs suggestion specific targeting of BRAF protein. Reduction of BRAF protein levels was noted also in wild type cells suggestive of high affinity of BRAF 600E point mutation mAb to also wild type allele.

**Figure 5.**
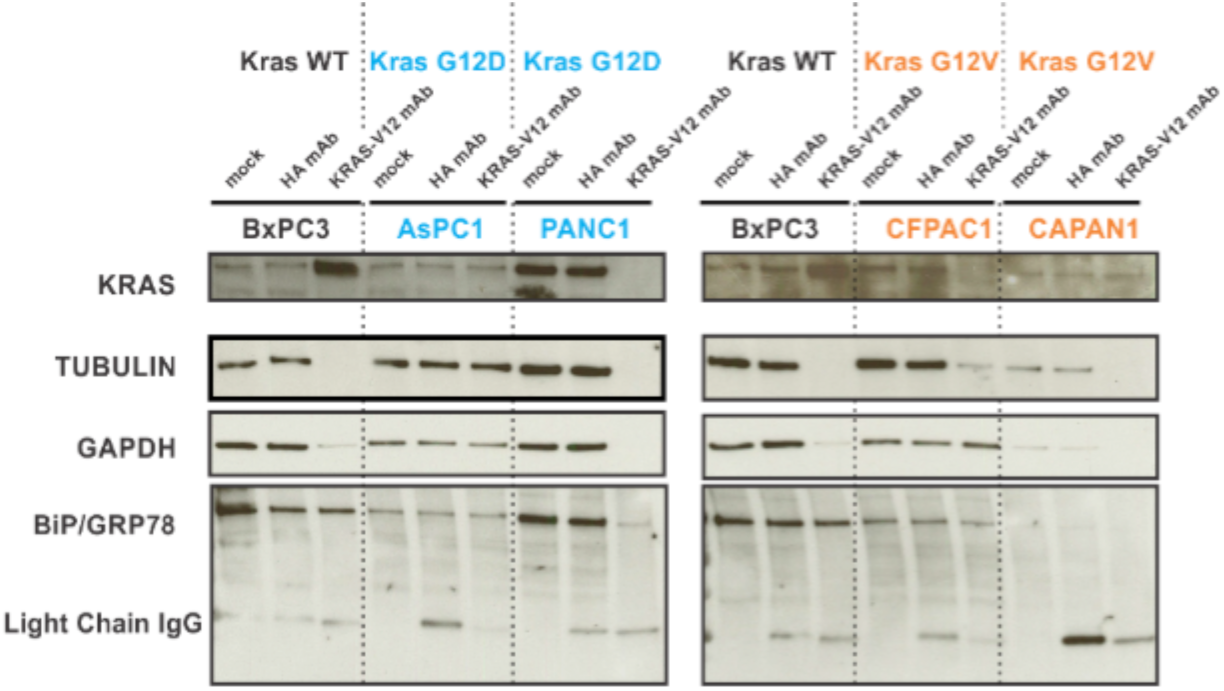
Precise depletion of *KRAS* somatic variant (KRAS p.G12V/D) in human pancreatic cancer cells. KRAS V12 point mutation specific mAbs (New East Biosciences, preservative-free formulation) was delivered into human pancreatic cell lines (BxPC3, AsPC-1, PANC-1, CFPAC-1, CAPAN-1) using ProJECT protein transfection reagent (Life Technologies) at 1:1 ratio. All cancer cell lines except BxPC3 (KRAS wild type) harbour the somatic mutation KRAS p.G12V or KRAS p.G12D. Two days after transfection cell lysates were analysed by immunoblot for expression levels of KRAS. Wild type KRAS expression was elevated in BxPC3 cell line whereas, mutated KRAS was specifically depleted in PANC-1 and CFPAC-1.

**Fig. 6.**
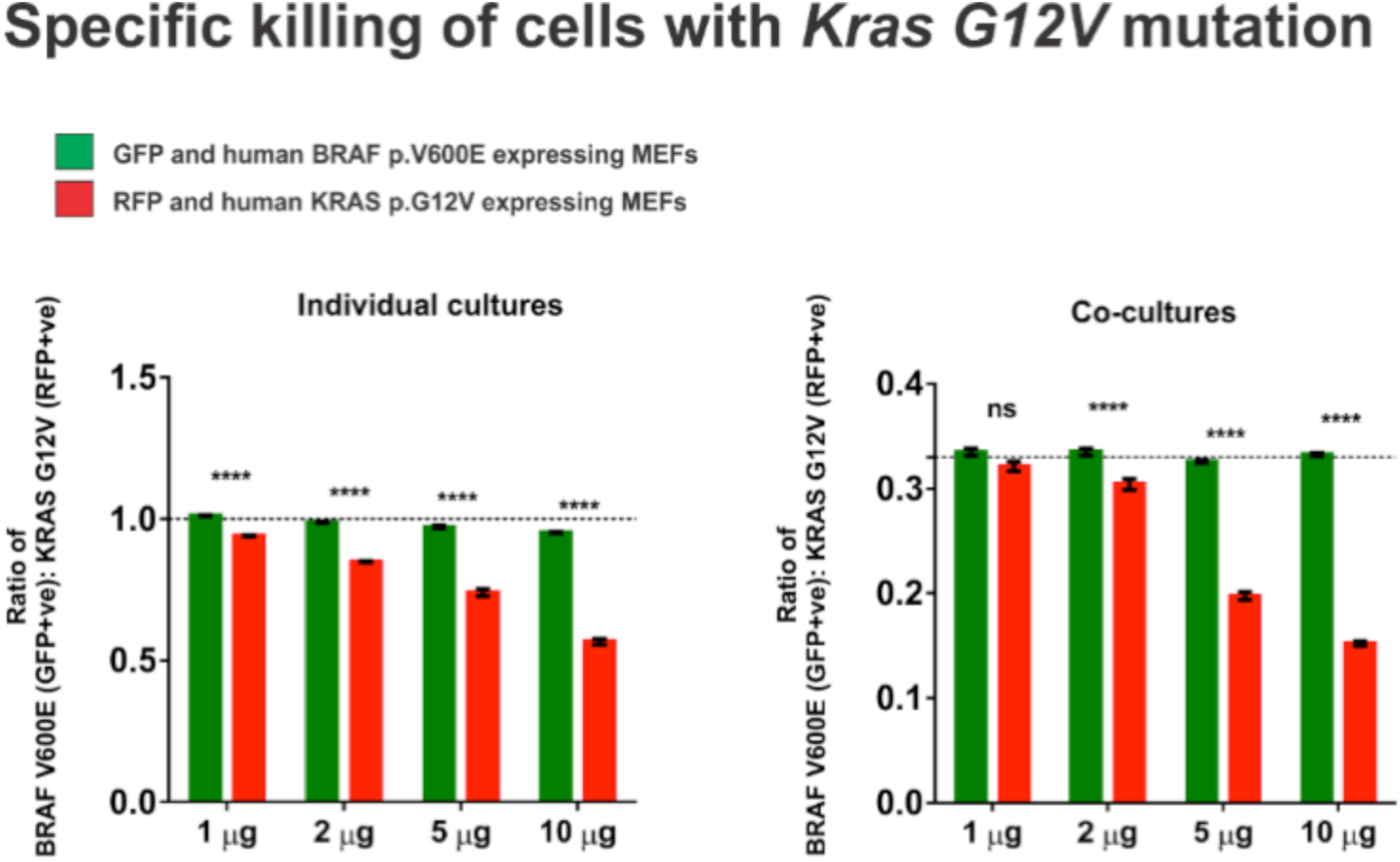
KRAS p.G12V somatic variant-specific inhibition of growth in co-cultivated cells. Mouse embryonic fibroblasts (MEFs) stably expressing somatic variants namely human BRAF p.V600E and human KRAS p.G12V were genetically labelled using GFP and RFP expressing constructs respectively. All exogenous expression was directed by CAG promoter and are stably integerated into the genome (transposon based plasmid construct). Indicated monoclonal mAbs were delivered (ProJECT protein transfection reagent (Life Technologies) at varied amounts and proportion of cells remaining were estimated 48 hours post treatement by flow cytometry by gating on beads. Shown are results from three independent experiments. Cell numbers are shown in percentages of total living cells.

**Fig. 7.**
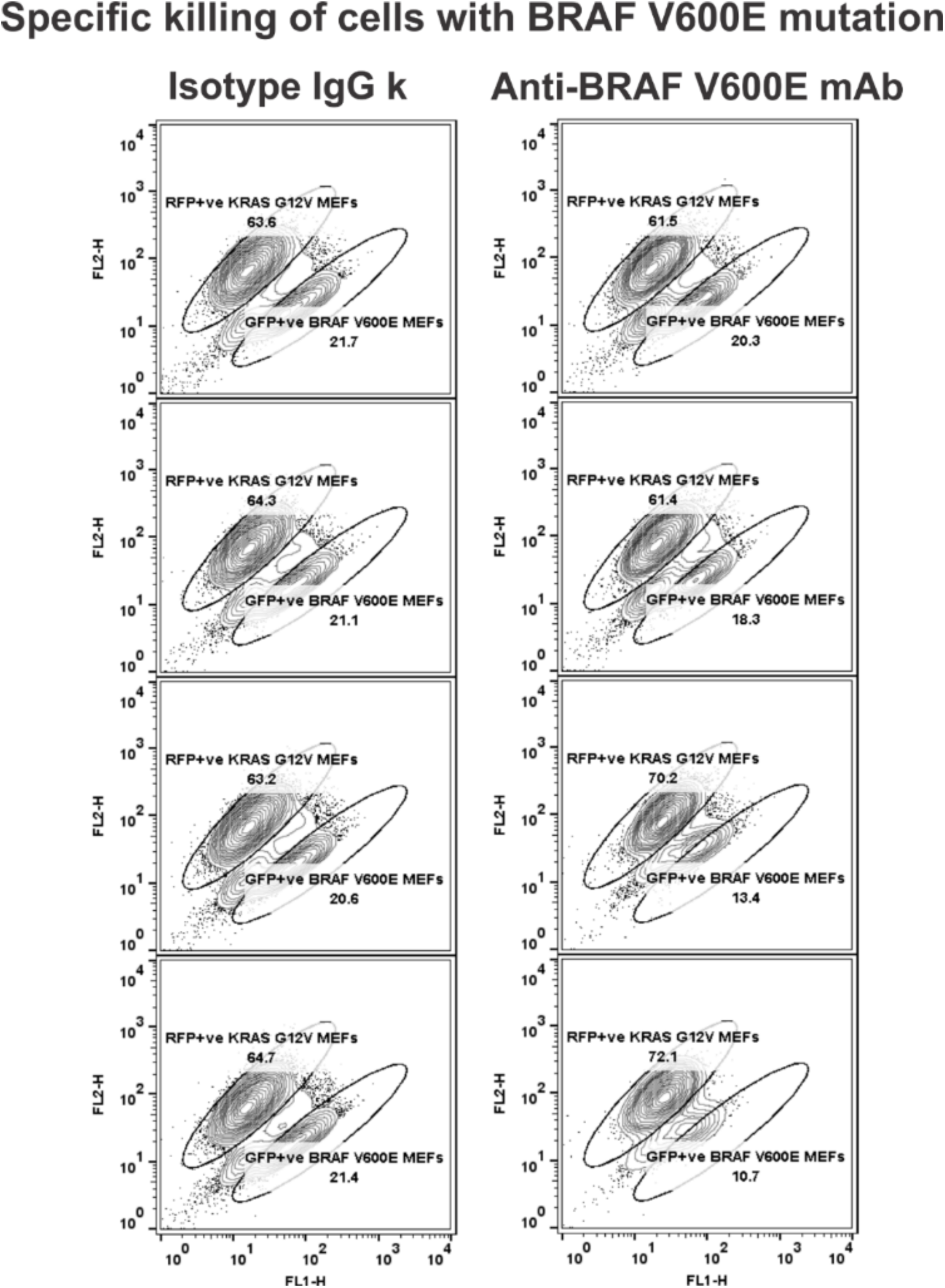
BRAF p.V600E somatic variant-specific inhibition of growth in co-cultivated cells. Mouse embryonic fibroblasts (MEFs) expressing mutated proteins namely human BRAF p.V600E (GFP+ve cells) and human KRAS p.G12V (RFP+ve cells) were co-cultivated under identical growth conditions. Monoclonal mAbs were delivered (ProJECT protein transfection reagent (Life Technologies)) at indicated amounts and proportion of cells remaining were estimated 48 hours post treatement by flow cytometry by gating on beads. Shown is the representative of three independent experiments. Cell numbers are shown in percentages of total living cells.

**Figure 8.**
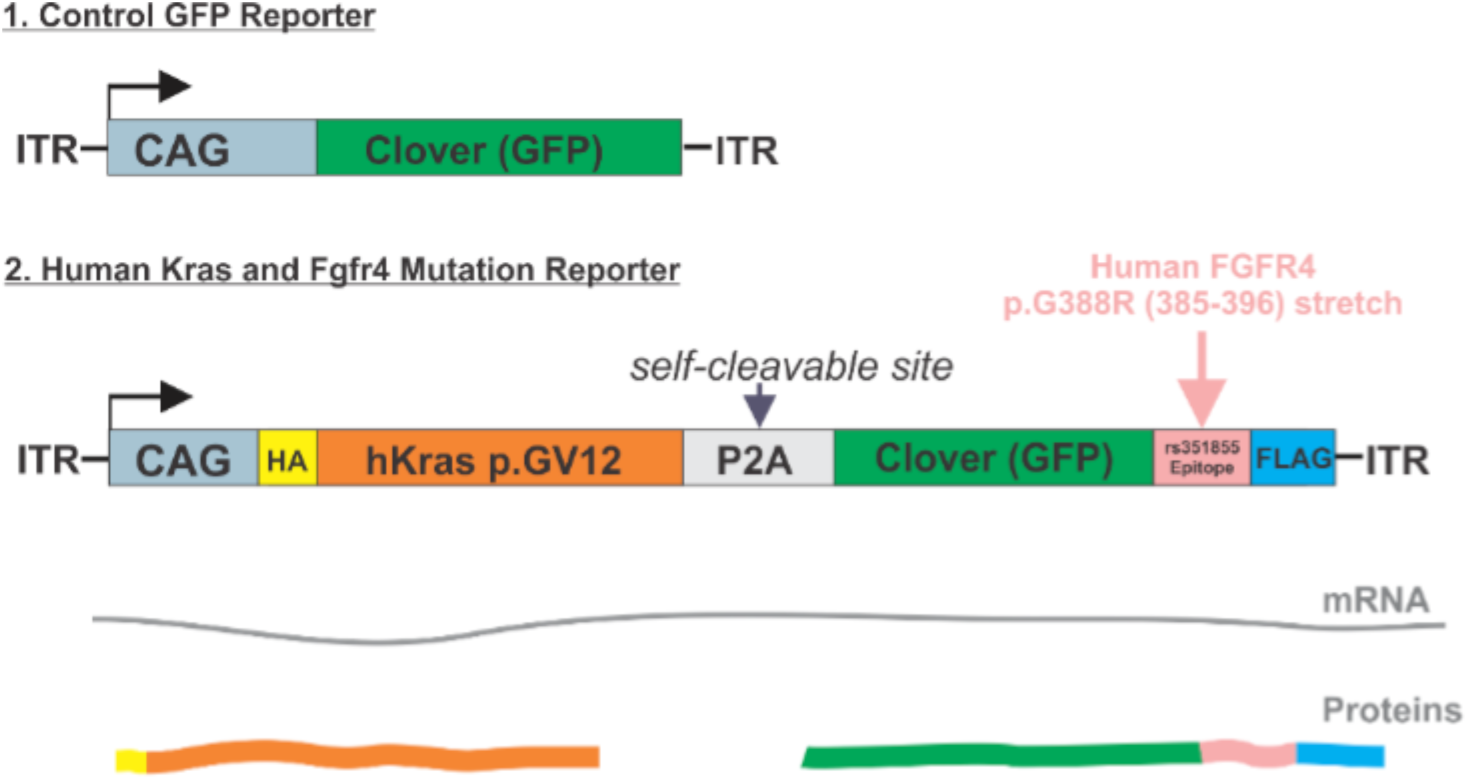
Generation of genetic reporters for screening and optimization protocols. Transposon based reporter expression plasmids shown above were constructed for generating stable cell lines. These cell lines were used for functional assays as well as for downstream protocols such as for screening assays, confocal live FRET imaging, mass spectrometry and co-culture assays. Importantly, multiple epitope proteins were expressed using self-cleavable peptide sequences to allow stoichiometric expression and to answer if inhibiting translation of one epitope interferes with others. This will be suggestive of mRNA knockdown.

**Figure 9.**
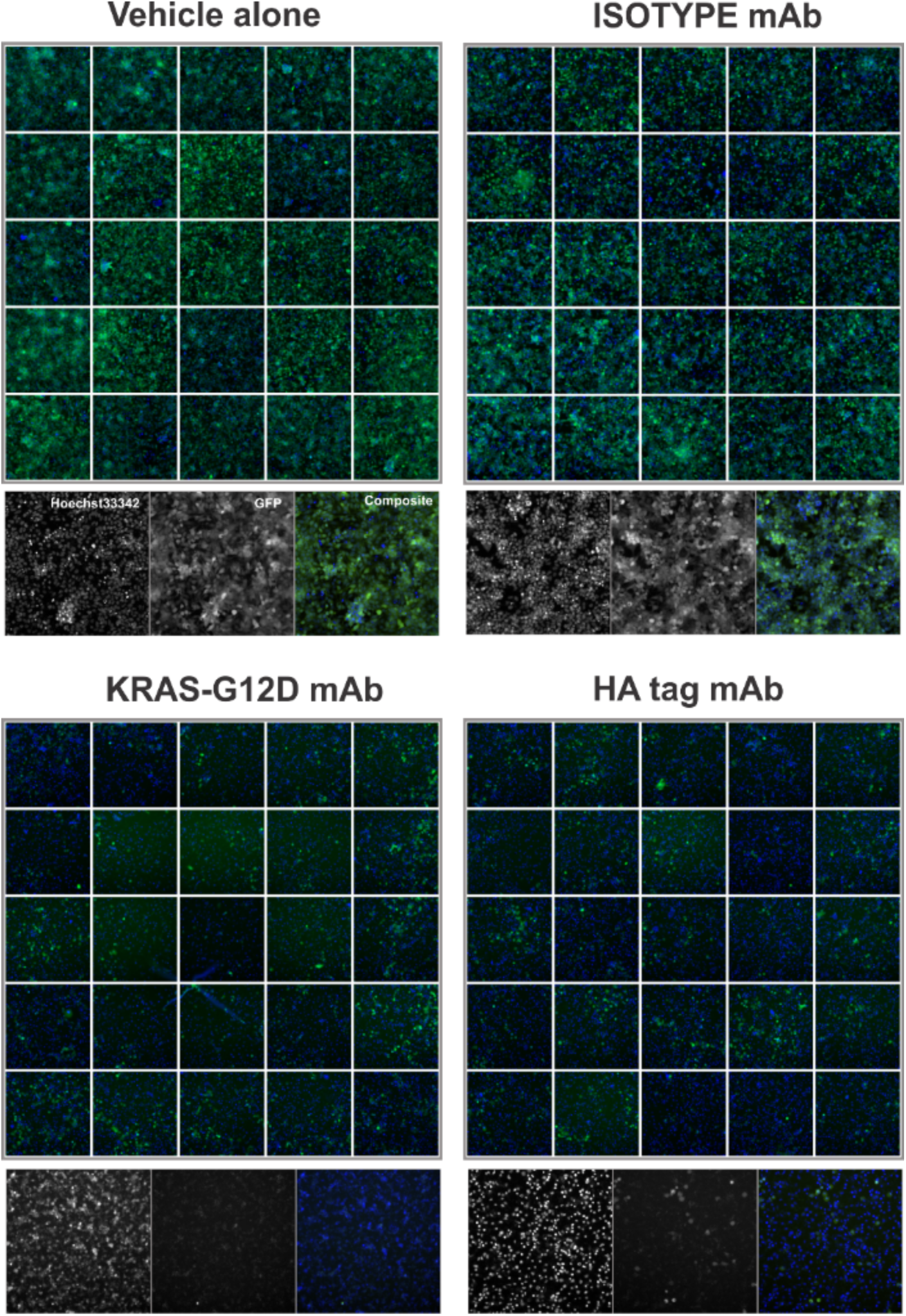
Inhibition of polycistronic target expression. Sorted HEK293T cells stably expressing HA-FGFR4 Arg388-GFP epitope reporter#2 (shown above) were transfected in replicates with indicated mAbs. Two days after treatment, cells were imaged at 10 X magnification for monitoring the target specific effects as indicated by GFP expression. Shown are representative images from one of three independent experiments. Green: GFP fluorescence and Blue: Nuclear staining by Hoechst 33342.

## Methods

### Cell lines and Medium

Mouse embryonic fibroblasts (MEFs) were generated from wild type and rs351855 SNP-knock in mice as described before (Ulaganathan et al, 2015)^1^. Human cancer cell lines used in this study were as follows NCI-H1944 (Lung), NCI-H1838 (Lung), BXPC3 (Pancreas), PANC1 (Pancreas), HACAT (Skin) and SK-MEL-3 (Skin). All cell lines used were obtained from American Type Culture Collection (ATCC) and authenticated in-house using a StemElite ID system (Promega, G9530). None of the cell lines used in this study were in the International Cell Line Authentication Committee list of currently known cross-contaminated or misidentified cell lines. Cell lines maintained by our cell bank staff are routinely controlled for mycoplasma contamination. The cell lines used in this study were confirmed to be free of any mycoplasma contamination.

### Plasmid Expression Constructs

Multi-cistronic transposon based plasmid constructs were generated by LRII clonase reaction between pENTRY-D TOPO entry vector and destination vector containing ITR-CAG-dest-IRES-puro-P2A- thy1.1-ITR. DNA encoding HA-tagged human KRAS p.G12V protein variant and FLAG-tagged EGFP protein fused to short antigenic epitope of FGFR4 p.G388R protein variant was cloned into pENTRY- D TOPO. To genetically label mouse embryonic fibroblasts (MEFs) with green fluorescent proteins (EGFP) and red fluorescent proteins (DsRED) transposon based plasmid DNA encoding EGFP and DsRED driven by the CAG promoter. Genetically labelled MEFs were selected by puromycin followed by flow sorting of DNA encoding EGFP and DsRED was used.

### Generation of stable cell lines

Stable cell lines were generated by co-transfection in 1:1 ratio of pCMV-SB transposase and ITR- plasmids and selected by puromycin resistance for a week followed by three rounds of flow cytometry sorting using BD FACSAriaIII.

### Immunoprecipitation

Stable engineered cell lines were lysed using RIPA buffer (#9806, Cell Signaling) and HA-tagged proteins and Venus fluorescent proteins were immunoprecipitated using HA-tag (C29F4) Rabbit mAb (Magnetic Bead Conjugate) (#11846, Cell Signaling) and GFP-Trap_M (#gtm-20, Chromotek) beads respectively. BCA-normalized lysates were incubated with magnetic beads at 4 degrees Celsius for 4 hours. Interacting proteins were isolated from the lyaste using magnets and washed with RIPA buffer three times.

### Western Blot analysis

Whole-cell lysates were prepared using 1× cell lysis buffer (Cell Signaling, 9803) containing cOmplete, mini, EDTA-free tablets (Roche 11836170001) and PhosSTOP tablets (Roche 04906837001). Equal concentrations (20–30 μ g) were loaded after a (BCA) assay, were run out on 4–15% Mini-PROTEAN TGX Gels (Bio-Rad 456-1083) and subsequently transferred onto a nitrocellulose membrane. The blots were blocked in 1× NET-Gelatin buffer (1.5 M NaCl, 0.05 M EDTA, 0.5 M Tris pH 7.5, 0.5% Triton X-100 and 0.25 g ml-1 gelatin) and incubated with primary antibodies overnight at 4 °C. Fractions of cell membranes were prepared using a FOCUS Membrane Protein Kit (G Biosciencs, 786-249); cytoplasm and nucleus were prepared using Nucbuster (Novagen, 71183-3).

For quantitative assessments of protein expression normalized lysates were analyzed by standard immunoblotting techniques using following antibodies: Phospho-Tyrosine (P-Tyr-1000) Multimab Rabbit mAb mix (HRP Conjugate) (#12695, Cell Signaling), HA-tag (C29F4) Rabbit mAb (HRP Conjugate) (#14031, Cell Signaling), GFP (D5.1) XP Rabbit mAb (HRP Conjugate) (#2037, Cell Signaling), Phospho-STAT3 (Tyr705) (D3A7) XP Rabbit mAb (#9145, Cell Signaling), STAT3 (124H6) Mouse mAb (#9139, Cell Signaling), α-Tubulin Mouse mAb (DM1A) (#T 9026, Sigma Aldrich), Homemade GFP Mouse mAb, FGFR4 (D3B12) XP Rabbit mAb (#8562, Cell Signaling), p44/42 MAPK (Erk1/2) (137F5) Rabbit mAb (#4695, Cell Signaling), phosphor-p44/42 MAPK (Erk1/2) (Thr202/Tyr204) (20G11) Rabbit mAb (#4376, Cell Signaling), EGFR (D38B1) XP Rabbit mAb (#4267, Cell Signaling), RAS (D2C1) Rabbit mAb (#8955, Cell Signaling), BRAF (D9T6S) Rabbit mAb (#14814, Cell Signaling), β-Actin (D6A8) Rabbit mAb (#8457, Cell Signaling), BiP/GRP78 Mouse mAb (40/BiP) (#610979, BD Transduction Labs), FGFR4 Rabbit pAb (C-16) (#sc-124, Santacruz Biotechnology) and Mouse anti Rabbit Igg (Conformation Specific) (L27A9) mAb (HRP Conjugate) (#5127, Cell Signaling)

### Intracellular-delivery of mAbs

Preservative-free formulations of mAbs in PBS were conjugated to cationic lipids using ProJECT (#89850, ThermoScientific) reagent as per manufacturer’s instruction manual and delivered to cells in serum-free cultivation medium. Two days post delivery, cells were washed and lysed in 1X cell lysis buffer (#9803, Cell Signaling). Mutation specific mAbs namely B-raf (V600E), Cat# 26039 and Ras (G12D), Cat# 26036 were obtained from NewEast Biosciences. FGFR4 pG388R variant specific mAbs reactive against mouse and human epitopes were custom made using services from AbPro Labs.

### Flow cytometry

GFP and RFP labelled cells and co-cultivated cells were analysed using FACS Calibur (BD Biosciences) in cold PBS.

### Fluorescence microscopy

Confocal imaging of GFP and RFP labelled cells was done using Lecia TCS SP8 microscope (Leica Microsystems).

Knock-down of GFP expression two days after intracellular delivery of lipid-conjugated antibodies, cells were imaged using Cellomics ArrayScan VTI HCS Reader (Thermo Scientific) using Hoescht filters and GFP filters.

## Acknowledgements

The authors thank Martin Spitaler, Markus Oster for assistance with confocal imaging and flow cytometry based sorting of genetically labelled cells. The authors thank Bianca Sperl for technical and secretarial assistance.

## Author contributions

VKU conceived the project, designed and performed the experiments, analysed the data, interpreted the results and wrote the manuscript. AU supported this work and reviewed the drafts of the manuscript.

